# Copy number alterations and epithelial-mesenchymal transition genes in diffuse and intestinal gastric cancers in Mexican patients

**DOI:** 10.1101/2021.11.22.469612

**Authors:** Violeta Larios-Serrato, José-Darío Martínez-Ezquerro, Hilda-Alicia Valdez-Salazar, Javier Torres, Margarita Camorlinga-Ponce, Patricia Piña-Sánchez, Martha-Eugenia Ruiz-Tachiquín

## Abstract

Gastric cancer (GC) is a malignancy with the highest mortality among diseases of the digestive system worldwide. The study of GC-alterations is crucial to understand tumor biology, to establish important aspects of cancer prognosis and treatment response. Here, we purified DNA and performed whole-genome analysis with high-density arrays in samples from Mexican patients diagnosed with GC: diffuse (DGC) or intestinal (IGC), or non-atrophic gastritis (NAG) samples that served as controls. We identified shared and unique copy number alterations (CNA) between these altered tissues involving key genes and signaling pathways associated with cancer, allowing their molecular distinction and identification of the most relevant molecular functions impacted. When focused on epithelial-mesenchymal transition (EMT) genes, our bioinformatic analysis revealed that the altered network associated with chromosomal alterations included 11 genes shared between DGC, IGC, and NAG, as well as 19 DGC- and 7 IGC-exclusive genes, whose main molecular functions included adhesion, angiogenesis, migration, metastasis, morphogenesis, proliferation, and survival. This study presents the first whole-genome high-density array study in GC from Mexican patients and reveals shared and exclusive CNA-genes in DGC and IGC. In addition, we provide a bioinformatically predicted network focused on CNA-altered genes involved in the EMT, associated with the hallmarks of cancer, as well as precancerous alterations that could lead to gastric cancer.

**Implications:** Molecular signatures of diffuse and intestinal GC, predicted bioinformatically, involve common and distinct CNA-EMT genes related to the hallmarks of cancer that are potential candidates for screening GC biomarkers, including early stages.

## Introduction

According to Global Cancer Observatory (GCO or GLOBOCAN) statistics, cancer is the leading cause of death in the World with 9.9 million deaths in 2020. This year, cancer had an incidence of 20% in the Caribbean and South America with high mortality rates (14%). Worldwide, gastric cancer (GC) is estimated to be the fifth most common cancer in both genders, ranking sixth in new cases with over one million per year and third in mortality (1). In Mexico, according to statistics from the Instituto Nacional de Estadística y Geografía (INEGI), three out of ten cancer deaths in the age group among 30-59 years old were due to cancer of the digestive system. From 2011 to 2016, four out of 10 and three out of 10 cancer deaths in women and men aged 60 years old or more, respectively, resulted from tumors in digestive organs (2).

GC refers to any malignancy originating in the region between the gastroesophageal junction and the pylorus. The World Health Organization and the Lauren classification system (3) have described two types of GC: intestinal and diffuse. The intestinal or differentiated gastric cancer (IGC) is characterized by localized and expansive growth, while diffuse gastric cancer (DGC) has an infiltrating growth pattern, is an undifferentiated adenocarcinoma, and presents dispersed cells with individual or group invasive capacity (4). The development of IGC is preceded by a precancerous process of several years and stages: active chronic gastritis, multifocal atrophic gastritis, complete intestinal metaplasia, incomplete intestinal metaplasia, dysplasia and adenocarcinoma (5). GC has a multifactorial origin: diet, lifestyle, genetics, socioeconomic factors, and it has been observed that 80% of cases of IGC are associated with previous *Helicobacter pylori* infection (6,7). GC is characterized by a complex etiology with a set of factors, such as genetic alterations and external factors. However, it has been reported that less than 3% of GC is due to heredity, and involves hereditary DGC, proximal polyposis of the stomach, and hereditary colorectal cancer not associated with polyposis (6). Regarding molecular pathogenesis, chromosomal instability (aneuploidy, chromosomal translocation, amplification, deletions and loss of heterozygosity), gene fusion and microsatellite instability (hypermethylation of gene repair promoters) are involved (7).

The number of copy alterations (CNA) represent a class of genetic variation that involve cumulative somatic variations, CNA are defined as non-inherited genetic alterations that occur in somatic cells (8). These unbalanced structural variants usually contain gains or losses. Its interpretation and the CNA report continue to be a topic of interest in health, and become evident to play an important role in GC (9,10).

The majority of gastric adenocarcinomas, like many other solid tumors, show defects in the maintenance of genome stability, resulting in DNA CNA that can be analyzed by comparative genomic hybridization (CGH) (11). There is a widespread and common phenomenon among humans and several studies have focused on understanding these genomic alterations that are responsible for cancer and might be used in diagnosis and prognosis (12).

Currently, there are few published studies involving genotyping of GC samples employing high-density microarrays (13–15); however, in those altered chromosomes, gains and losses have a phenotypic impact, and different signaling pathways are involved. The presence of CNA changes the genetic dose and would modify several molecular mechanisms as epithelial-mesenchymal transition (EMT), which is the transformation of epithelial to mesenchymal cells and a critical stage for the transition to metastasis (16). Currently, there are more than 1184 genes at the Epithelial-Mesenchymal Transition Gene Database 2.0 (dbEMT 2.0), which are involved in other cancer-related processes such as proliferative signaling, evading growth suppressors, avoiding immune destruction, disabling replicative immortality, tumor-promoting inflammation, inducing angiogenesis, genome instability, mutation, resting cell death, deregulation cellular energetic activity, invasion and cell plasticity (17). EMT includes activation of transcription factors, expression of specific cell-surface proteins, reorganization and expression of cytoskeletal proteins, and production of extracellular matrix (ECM) degrading enzymes (18). EMT has been linked to the progression of cancer and increased stemness of tumors (18,19), and observed in the formation, invasion and metastasis of GC (20,21). Also, a relationship has been established between the presence of CNA and its effect on the expression of EMT genes in different cancer cell lines (22). Reports of CNA events involving Latin American populations are still scant (23,24). In fact, at present there are few studies of GC genotyping by whole-genome high-density microarrays in Mexican patients (14,15). Therefore, our study aimed to determine CNA in DGC and IGC to identify through bioinformatics analyses, the main genes and signaling pathways involving EMT-genes.

## Material and Methods

### Samples

Institutional Review Board approval was obtained for the study origin (Register number: 2008-785-001). Clinical data and patient samples were processed under informed consent. All samples were collected over three years (April 2010-May 2013) following standardized endoscopy preservation protocols (25). The histology of the biopsies was assessed independently by two trained pathologists. They assigned the phenotypic diagnosis of diffuse or intestinal tumors and non-atrophic gastritis (NAG) samples. Only samples with identical results were included in the study.

We included 21 patients (5 females and 16 males) with tissue samples that met the quality criteria from DGC (n=7) and IGC (n=7) diagnoses, as well as NAG (n=7) as controls. To identify the most relevant alterations for GC, we focused our analysis on alterations present in at least three patients as an arbitrarily threshold (cut-off ≥ 3 patients; ≥ 40% samples).

### DNA extraction

DNA extraction was done with a commercial kit (QIAamp^®^ micro Kit, QIAGEN^®^) according to the manufacturer’s instructions. The extraction was modified to include an initial incubation at 95°C for 15 min followed by 5 min at room temperature as described previously, before being digested with proteinase K for three days at 56°C in a water-bath adding fresh enzyme at 24 h intervals (26).

### DNA quality assessment and preparation

The extracted DNA was quantified by spectrophotometry (Nanodrop 2000, Thermo Scientific). Multiplex PCR was done to assess the quality of DNA (Multiplex PCR kit, QIAGEN^®^) with a set of primers to amplify various regions of the GAPDH gene (27). Products were visualized by electrophoresis (RedGel^®^ Nucleic acid gel stain, Biotium) on a 1% agarose gel and documented under an ultraviolet light transilluminator system (Syngene, Frederick, MD, USA).

### High-density whole-genome microarray analysis

Samples were analyzed by Affymetrix^®^ CytoScan™ microarrays according to the manufacturer’s protocol beginning with 250 ng DNA, except for the addition of five PCR cycles to increase the DNA sample. PCR products (90 micrograms) were fragmented and labeled using the additional PCR.

### Copy number processing

Raw intensity files (.CEL) retrieved from the commercial platform were analyzed using their proprietary software Chromosome Analysis Suite v3.2, using NetAff 33 Libraries based on the construction of the hg19 genome (Feb 2009) as a reference model.

Data processing was based on the segmentation algorithm, where the Log2 ratio for each marker was calculated relative to the reference signal profile. To calculate CNV, the data were normalized to baseline reference intensities using the reference model (provided by ChAS) including 270 HapMap samples, as well as 96 healthy normal individuals. The Hidden Markov Model (HMM) available in ChAS was used to determine the copy number state (CN-state) and their breakpoints. The customized high-resolution condition was used as a filter for the determination of CNV: CN-gains with 50 marker count and 400 Kb, and CN-losses with 50 marker count and 100 Kb. We used the median absolute pairwise difference (MAPD) and the single nucleotide polymorphism quality control (SNP-QC) score as the quality control parameters. Only samples with values of MAPD > 0.25 and SNP-QC < 15 were included in further analysis.

### Bioinformatic analysis

We developed a Perl script to load the CNV segment data files generated by ChAS for each sample, compare the files to build a table of genes that contains events types (gains or losses), frequencies of altered regions, including chromosomes and cytogenetic bands, and Online Mendelian Inheritance in Man (OMIM) information, and incorporate additional information from different databases: haploinsufficiency information from DECIPHER database of genomic variation, genes reported at dbEMT 2.0, and genes affected in gastric adenocarcinoma from Harmonized Cancer Datasets (Table S1).

The genes altered in at least three patients (cut-off ≥ 3) with DGC, IGC or NAG were included for analysis and visualizations with the language and environment for statistical computing and graphics R v4.0.2 and Bioconductor v3.12 packages.

The karyotype was created with KaryoploteR and BSgenome.Hsapiens.UCSC.hg19 v1.4.0. The comparison among samples was made using Venn diagrams with the jVenn server and the heatmap with gplots. Enrichment and gene ontology (GO) analysis were performed with the ClusterProfiler v3.16.1 packages, org.Hs.eg.db 3.11.4, enrich plot v1.8.1, and GOplot v1.0.2, with the support of functional enrichment analysis via DAVID v6.8 bioinformatics resources. The profile of altered molecular functions in GC was summarized according to the proportion of CNA-genes and the Molecular Function (MF) gene collection of the gene ontology terms from the DAVID database, adjusted by FDR. We used dot plots, heatmaps, and chord plots to visualize the general GC CNA profiles for DGC, IGC, and NAG.

To identify the main genes and signaling pathways involving CNA EMT-genes, we analyzed and compared GC CNA-genes (cut-off ≥ 3) according to those previously reported in the dbEMT 2.0 accessed on October 12^th^, 2020.

Finally, to establish the profile-associated hallmarks of cancer involving DGC, IGC, and NAG EMT-genes, we generated an interaction network by CNA-type (gains and losses) based on genetic and physical interactions, biological pathways, and predicted relationships using the GeneMANIA prediction server and Cytoscape v.3.8.2, including the manual annotation of their corresponding cancer hallmarks: adhesion, angiogenesis, inflammation, migration, metastasis, morphogenesis, proliferation, and survival (28) with punctual scrutiny and help from databases such as The Human Protein Atlas. For further consultation of the databases, protocols, software, and specific packages used in this study see Table S2.

## Results

### Sample characteristics

Samples from 21 Mexican patients (third-generation Mexicans) between 35 and 91 years old (61.7 ± 15.9 years) without previous cancer treatment (naïve) were included in our study. The data includes 21 samples: seven DGC, seven IGC and seven NAG (control samples). Our data have been deposited in the NCBI Gene Expression Omnibus and assigned the GEO series accession number GSE117093. We have seven adjacent tissue files (.CEL), but these files were not included in data analysis, because some adjacent tissues were contaminated with cancer cells or these were not of the quality required for subsequent analyzes.

Table 1 shows the identity card (ID) and the percentage of neoplastic cells for tumor tissues ranging between 50-70%. Blood agar culture showed that one IGC and three NAG patients were positive for *Helicobacter pylori* (data obtained from our biobank database). Tumor size (T), number of lymph nodes (N) and metastasis (M) classification data are also presented in Table 1.

**Table 1.** Characteristics of GC samples analyzed in this study.

| **ID** | **Age**<br>**(years)** | **Gender** | **CT** | **% CC** | ***H. pylori*** | **TNM** | **Treatment** |
| --- | --- | --- | --- | --- | --- | --- | --- |
| 3CG-008 | 72 | M | Intestinal | 70 | Positive | IB T1 N1 M0 | Naïve |
| 3CG-126 | 80 | M | Intestinal | 60 | Negative | IIA T4 N0 M0 | Naïve |
| 3CG-128 | 91 | M | Intestinal | 70 | Negative | IIA T3 N2 M0 | Naïve |
| 3CG-046 | 52 | F | Intestinal | 60 | Negative | IV T4 N2 M0 | Naïve |
| 3CG-099 | 59 | M | Intestinal | 50 | Negative | II T3 N0 M0 | Naïve |
| 3CG-146 | 71 | M | Intestinal | 60 | Negative | IIB T3 N2 M0 | Naïve |
| 3CG-104 | 69 | M | Intestinal | 60 | Negative | II A T4 N0 M0 | Naïve |
| 3CG-047 | 58 | M | Diffuse | 70 | Negative | IV T4 N3 M0 | Naïve |
| 3CG-173 | 76 | M | Diffuse | 70 | Negative | II A T2 N3 M0 | Naïve |
| 8CG-004 | 76 | M | Diffuse | 70 | Negative | II T1 N0 M0 | Naïve |
| 1CG-001 | 45 | M | Diffuse | 60 | Negative | IV T4N2M1 | Naïve |
| 3CG-035 | 55 | M | Diffuse | 60 | Negative | IV T4N2M0 | Naïve |
| 3CG-042 | 64 | M | Diffuse | 50 | Negative | IV T4, N2 M0 | Naïve |
| 3CG-064 | 38 | M | Diffuse | 50 | Negative | IV T4 N2 M0 | Naïve |
| 4GB-001 | 64 | M | Gastritis | 0 | Negative | NA | NA |
| 4GB-031 | 62 | M | Gastritis | 0 | Negative | NA | NA |
| 4GB-015 | 35 | F | Gastritis | 0 | Negative | NA | NA |
| 4GB-025 | 39 | M | Gastritis | 0 | Positive | NA | NA |
| 4GB-033 | 76 | F | Gastritis | 0 | Positive | NA | NA |
| 4GB-036 | 38 | F | Gastritis | 0 | Positive | NA | NA |
| 4GB-042 | 77 | F | Gastritis | 0 | Negative | NA | NA |

ID, identification code; CT, cancer type; CC, cancer cells; M, male; F, female; NA, not applicable; TNM, T: the extent of the primary tumor; N: the absence or presence and extent of regional lymph node; M: the absence or presence of distant metastasis (29–31).

### Genomic detection of CNA

We obtained the total number of CNA and classified them as gains or losses for each chromosome in GC and NAG samples. By total CNA, DGC had more CNA than IGC (3505 and 2781, respectively), while there were 828 events in NAG samples. By tissue, we observed more gains than losses (G/L) in both cancer types, DGC (2310/1195) and IGC (1550/1231), but the opposite occurred in NAG (375/453) (Table S3).

To identify the most relevant CNA in GC and NAG, we analyzed alterations occurring in at least three patients (cut-off ≥ 3). This comparison showed a similar pattern for total CNA with more events in DGC (710), IGC (590), and NAG (332). Also, we observed more gains than losses (G/L) in DGC (516/194), IGC (314/276) and even in NAG (196/136), unlike when all patients were included. In addition, DGC had the highest number of gains and IGC had the highest number of losses (Table S3). In Table 2 we show chromosomes and sizes with gain and loss numbers, representative and summarized.

**Table 2.** Principal affected chromosomes by CNA cumulative length in diffuse gastric cancer, intestinal gastric cancer and non-atrophic gastritis.

| **Type** | **Chromosome** | **Gains** | **Losses** | **Length**<br>**(Mb-cl)** |
| --- | --- | --- | --- | --- |
| **DGC** | 1 | 327 | ----- | 117.9 |
|  | 4 | ----- | 155 | 40.8 |
|  | 5 | ----- | 148 | 74.23 |
| **IGC** | 1 | ----- | 148 | 33.78 |
|  | 8 | 365 | ----- | 139.8 |
|  | X | ----- | 66 | 167.1 |
| **NAG** | 6 | ----- | 28 | 0.40 |
|  | 7 | 21 | ----- | 0.20 |
|  | 14 | 10 | ----- | 3.02 |
|  | 17 | ----- | 15 | 1.86 |
|  | X | ----- | 87 | 0.47 |
|  | X | 207 | ----- | 1.20 |

DGC: Diffuse gastric cancer; IGC: Intestinal gastric cancer; NAG: Non-atrophic gastritis; Mb-cl: Megabases cumulative length.

To visualize the distribution of DGC and IGC chromosome gains and losses, we plotted the identified CNA present in a karyogram (cut-off ≥ 3), locating alterations according to the coordinates of the Human genome hg19 (Figure 1). The top five altered cytobands are shown in Tables 3 and S4.

**Figure 1.**
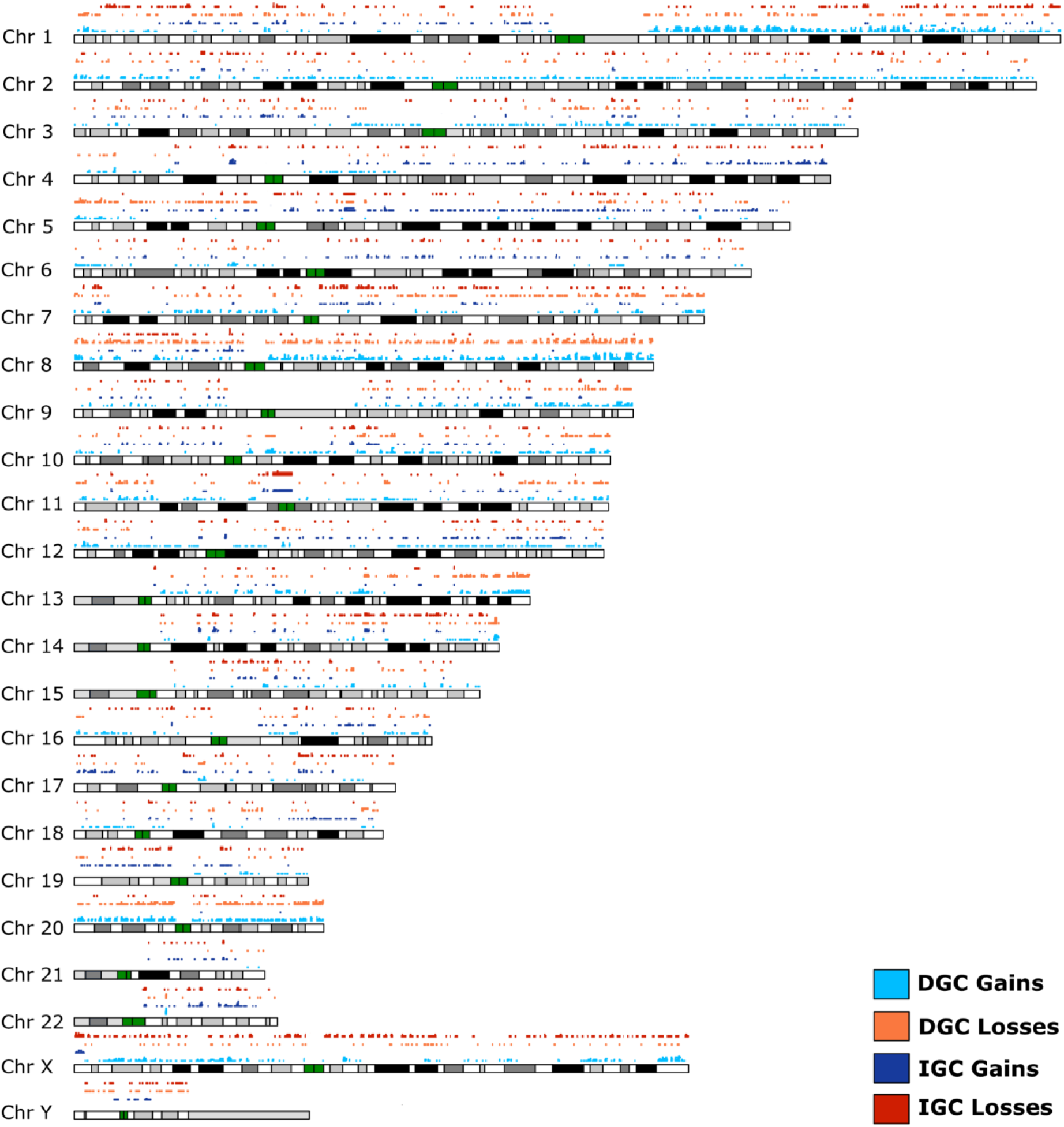
Karyogram with CNA distribution in diffuse and intestinal gastric cancers. CNA events (gains or losses) were present from chromosomes 1 to 22, X, and Y. Gains (blue and dark blue) and losses (orange and red) are plotted for ≥ 3 DGC or IGC patient samples. Cytobands (gray, black or white bars) and centromeres (green bars) are shown.

**Table 3.**
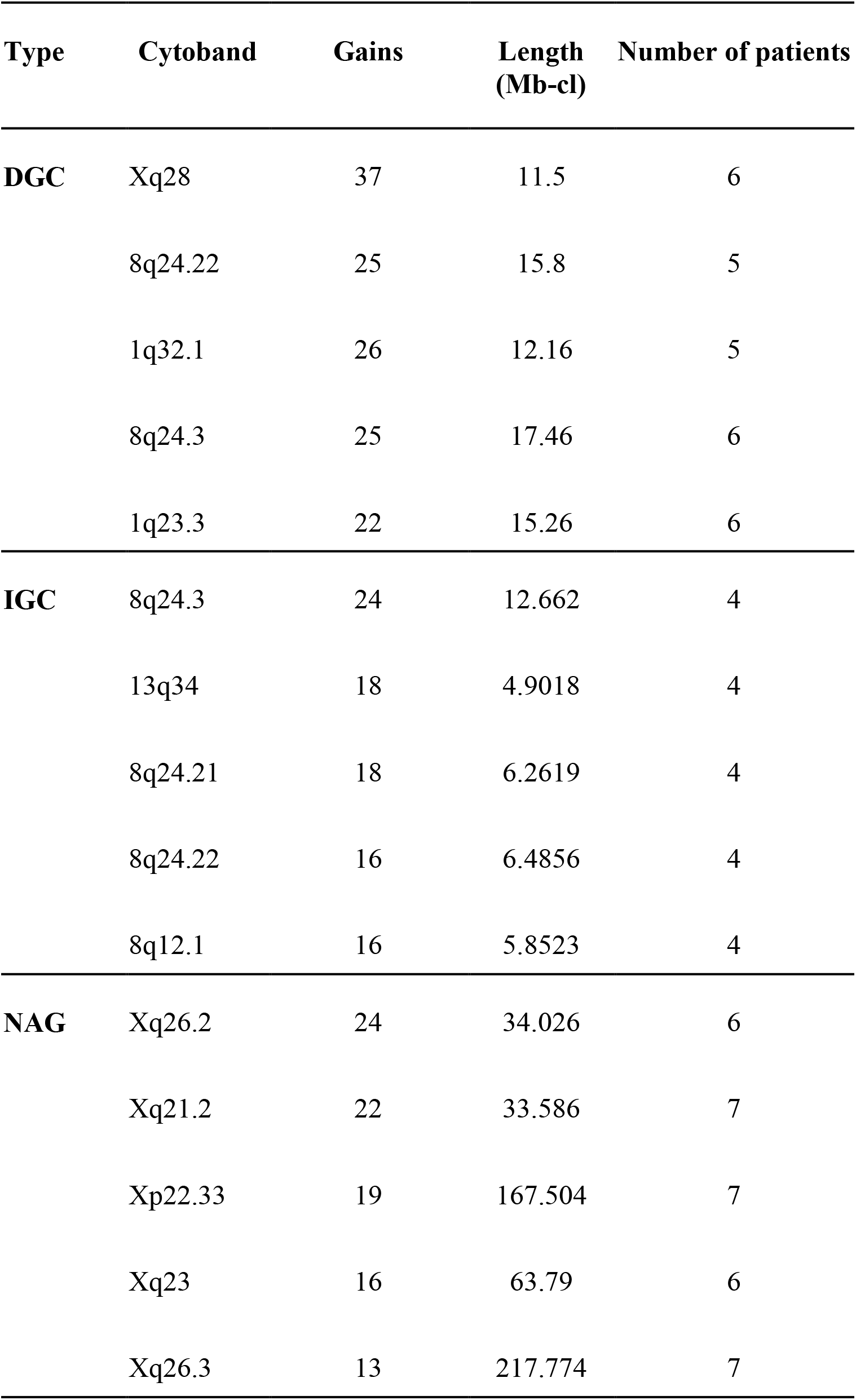
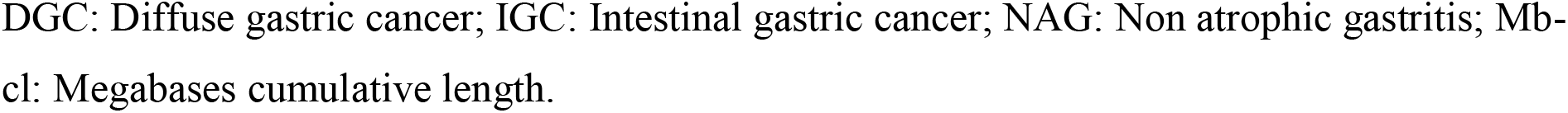
Top five altered cytobands in diffuse gastric cancer, intestinal gastric cancer and non-atrophic gastritis.

| Type | Cytoband | Gains | Length (Mb-cl) | Number of patients |
| --- | --- | --- | --- | --- |
| **DGC** | Xq28 | 37 | 11.5 | 6 |
|  | 8q24.22 | 25 | 15.8 | 5 |
|  | 1q32.1 | 26 | 12.16 | 5 |
|  | 8q24.3 | 25 | 17.46 | 6 |
|  | 1q23.3 | 22 | 15.26 | 6 |
| **IGC** | 8q24.3 | 24 | 12.662 | 4 |
|  | 13q34 | 18 | 4.9018 | 4 |
|  | 8q24.21 | 18 | 6.2619 | 4 |
|  | 8q24.22 | 16 | 6.4856 | 4 |
|  | 8q12.1 | 16 | 5.8523 | 4 |
| **NAG** | Xq26.2 | 24 | 34.026 | 6 |
|  | Xq21.2 | 22 | 33.586 | 7 |
|  | Xp22.33 | 19 | 167.504 | 7 |
|  | Xq23 | 16 | 63.79 | 6 |
|  | Xq26.3 | 13 | 217.774 | 7 |

DGC: Diffuse gastric cancer; IGC: Intestinal gastric cancer; NAG: Non atrophic gastritis; Mb-cl: Megabases cumulative length.

Interestingly, in DGC and IGC the most frequent CNA lengths were 100-200 Kb, while 1-50 Kb were more common in NAG, considering gains and losses (Table S5).

### Gastric cancer genes associated with CNA

Overall, we found 2441 CNA-genes in DGC-IGC-NAG. GC had 2420 affected genes (99%) while only 108 genes (4%) were affected in NAG; some of these alterations were shared between GC and NAG. We observed 1317 unique CNA-genes in DGC, 596 in IGC, and 21 in NAG; both cancer types shared 420 genes, while 60 genes were shared between GC and NAG. In addition, 19 NAG genes were shared with DGC and eight genes with IGC (Figure 2A and Table S6).

**Figure 2.**
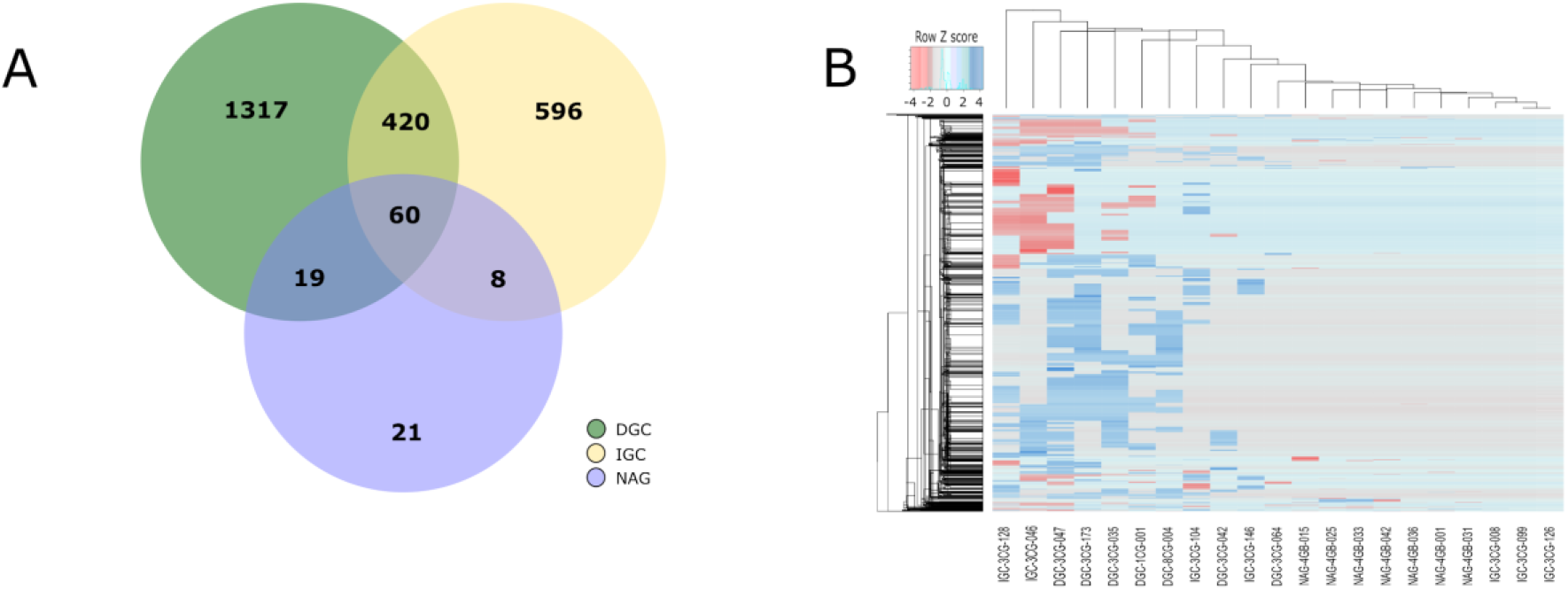
Profile of CNA-genes in gastric cancer from ≥ 3 patients. (A) The Venn diagram presents frequencies of specific and shared genes in diffuse gastric cancer (DGC), intestinal gastric cancer (IGC) or non-atrophic gastritis (NAG) in CNA. (B) The heatmap shows the hierarchical clustering of gains (blue), losses (red), and no alterations (light blue and light red).

To identify possible emerging patterns among samples, we performed hierarchical clustering heatmaps (Figure 2B). Results showed the molecular signature and hierarchical clustering of samples according to 2441 genes. The emerging pattern of altered genes affected by CNA distinguishes DGC and IGC from NAG.

### GO analysis of gastric cancer

We obtained the functional profile for GC through enrichment and gene ontology (GO) analysis of 1317 genes altered only in DGC and 596 in IGC. To identify the principal molecular functions altered in GC, we categorized these CNA-genes, independently of GC-type, in two groups: gains and losses (Figure 3A and 3B). The top ten molecular functions (MF) associated with CNA gains or losses revealed that transcription activator, tyrosine kinase activity, growth factors and hormone binding, as well as intracellular signal transduction genes were enriched in GC. Gene-losses mainly involved transcription coactivator and serine/threonine kinase activity, as well as, several receptors binding to hormone, steroid hormone, nuclear receptor, beta-catenin, intermediate filament, mitogen-activated protein kinase binding genes. In addition, we identified the principal CNA-genes affecting the MF by GC type: DGC and IGC (Figure 3C, 3D, and 3E).

**Figure 3.**
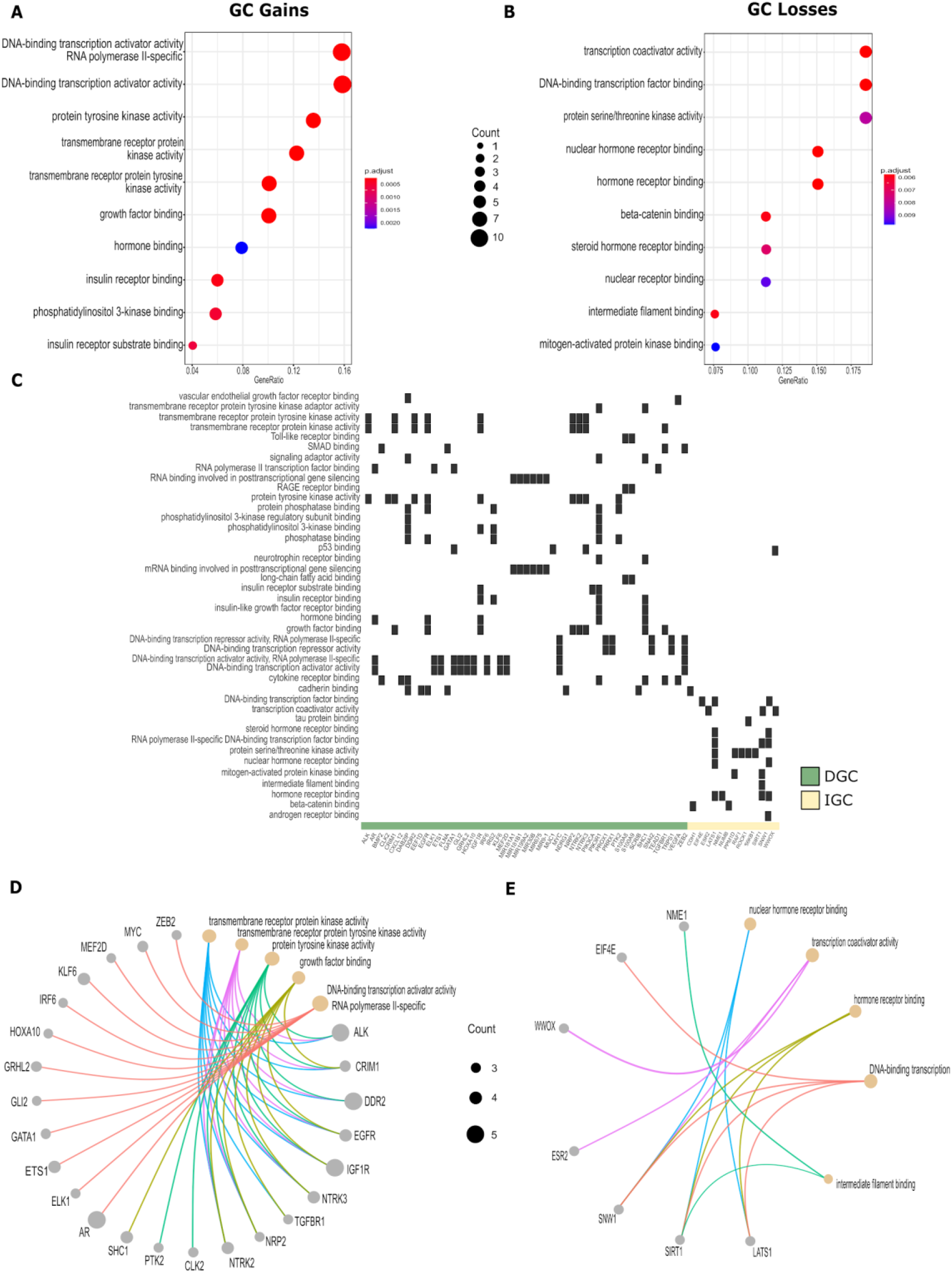
Molecular function profiles of gastric cancer CNA-genes. Gene enrichment analysis affected by CNA-genes in ≥ 3 gastric cancer (GC) patients. It is summarized by CNA-type with dot plots for gains (A) and losses (B), according to the proportion of CNA-genes and Molecular Functions (MF) gene collection of the gene ontology (GO) terms from the DAVID database, adjusted by FDR. The heatmap (C) shows the main MF altered by CNA-genes in Diffuse-GC and Intestinal-GC. The CNA-genes-MF networks depict the possible relationship between CNA gains (D) and losses (E) within GC. Gene-ratio (M/N) is the proportion between CNA genes in ≥ 3 GC patients (M) and the collection of genes from the GO term database function (N). Count (circle sizes) represents the number of CNA-genes associated with MF.

### CNA-EMT genes in DGC and IGC

To identify the main genes and signaling pathways involving CNA-EMT genes in GC and NAG, we compared GC CNA-genes against a comprehensive and annotated database of epithelial-mesenchymal transition genes (dbEMT). We found 551 CNA-EMT genes for DGC, 619 for IGC and 28 for NAG. With a cut off ≥ 3 in DGC-112, IGC-66 and NAG-5. The complete data of EMT-genes for DGC, IGC, and NAG with chromosome and cytoband locations, CNA-type (gain or loss), and *p*-values can be found in Table S1.

### GO Analysis of EMT-genes

We performed a gene ontology (GO) enrichment analysis to determine the MF of the main CNA-EMT genes affected in DGC, IGC, and NAG (Figure 4). Our analysis shows that CNA-EMT gene-gains in DGC involved transmembrane receptor tyrosine kinase, DNA and RNA binding, several receptor binding: insulin, growth factor, toll-like, hormone, as well as SMAD, p53, chromatin, calcium ion binding and microtubule binding (Figure 4A). CNA-EMT gene-losses included DNA and chromatin binding, nuclear hormone receptor binding, beta-catenin, steroid hormone, mitogen-activated protein binding, intermediate filament binding, p53 binding, RNA polymerase II-specific DNA binding (Figure 4B). Moreover, CNA-EMT gene-gains in IGC involved insulin receptor substrate and phosphatase binding, kinase regulation, neurotrophin receptor binding, 1-phosphatidylinositol-3-kinase activity, transmembrane receptor protein tyrosine kinase adaptor activity, and VEGF receptor binding; while CNA-EMT gene-losses in IGC involved only coenzyme binding and transcription coactivator activity (Figure 4C). On the other hand, the main MF for CNA-EMT gene-gains in NAG included transcription regulatory region DNA, transcriptional activator activity RNA, armadillo repeat and C2H2 zinc finger domain binding, gamma- and beta- catenin binding, as well as cysteine-type endopeptidase inhibitor activity involved in apoptotic process and estrogen receptor and steroid hormone receptor activity; while CNA-EMT gene-losses in NAG involved damaged DNA, WW domain, and p53 binding as well as DNA-binding transcription activator activity (Figure 4D).

**Figure 4.**
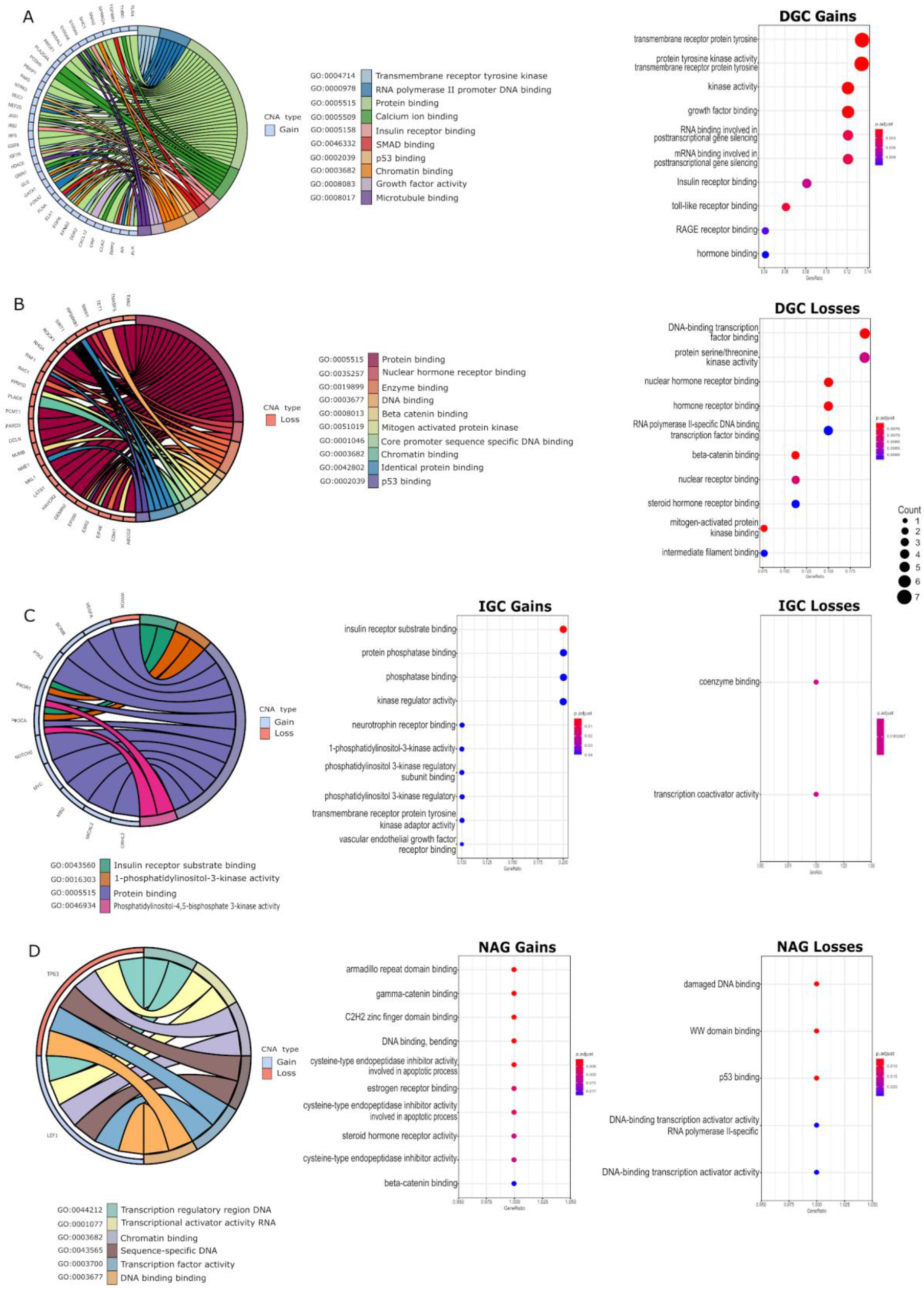
Principal molecular functions associated with CNA-EMT genes in gastric cancer and non-atrophic gastritis. The main MF associated with CNA-EMT are represented with chord plots and dot graphics for (A) DGC gains, (B) DGC losses, (C) IGC gains or losses, and (D) NAG gains or losses. Chord plots (left panels) show associations between genes and molecular functions, indicating their CNA-type by color coding (gains/blue and losses/red). Dot plots (right panels) show an enrichment analysis of molecular functions, and loss or gain genes counts in samples. CNA, copy number alteration; EMT, epithelial mesenchymal transition; DGC, diffuse gastric cancer; IGC, intestinal gastric cancer; NAG, non-atrophic gastritis.

### CNA-EMT genes associated hallmarks of cancer

Based on the main molecular profile of altered CNA-EMT genes in GC and NAG, we generated the functional network between 39 previously selected unique CNA-EMT genes, (19 genes for DGC, seven for IGC, 11 common to GC, and two for NAG; cut-off ≥ 3 patients). Gained genes with the highest degree, at least four interactions per gene, were EGFR, MICAL2, MYC, NDRG1, and PIK3R1, while lost genes included GLI2, EP300 and PTPN11. The principal functions related to these CNA-EMT genes have been previously associated with several hallmarks of cancer: adhesion, angiogenesis, inflammation, migration, metastasis, morphogenesis, proliferation, and survival (Figure 5).

**Figure 5.**
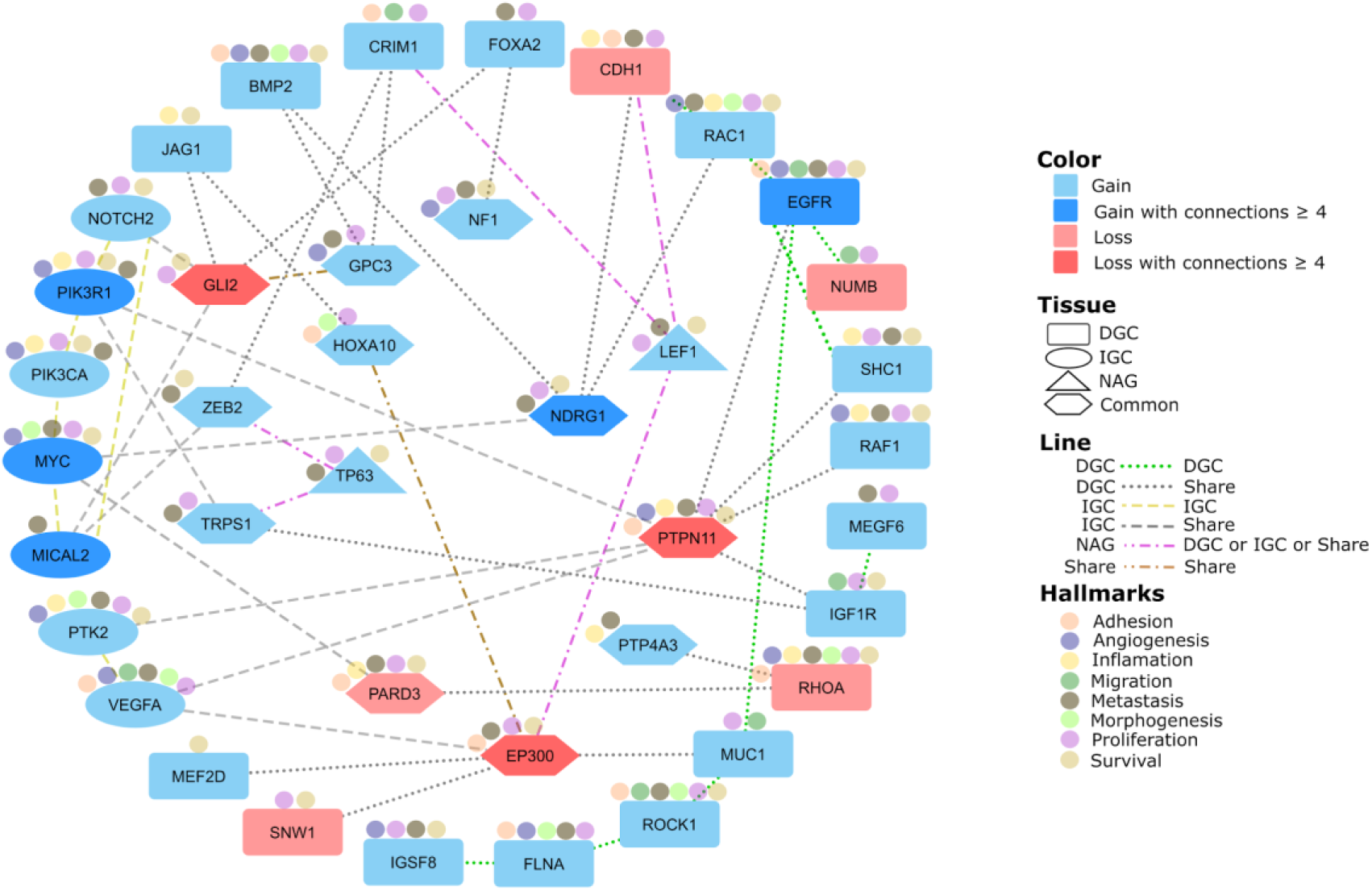
Gastric cancer and non-atrophic gastritis CNA-EMT genes network and associated hallmarks of cancer. The functional interactions among CNA-EMT genes identified in DGC (rectangles), IGC (ovals), and shared genes (hexagons) are identified by their CNA-type (gains/blue and losses/red) and associated hallmarks of cancer (colored dots). DGC, diffuse gastric cancer; IGC, intestinal gastric cancer; NAG, non-atrophic gastritis; EMT, epithelial-mesenchymal transition; CNA, copy number alterations.

## Discussion

This is the first whole-genome high-density array study in GC from Mexican patients in three groups: DGC, IGC, and NAG as non-cancerous control. Using this experimental strategy, it was possible to generate a karyogram and obtain molecular signatures for diffuse and intestinal GC, and their relationship with CNA-EMT genes independently of age, gender, % CC, *H. pylori* presence/absence, TNM, and treatment (naïve samples in our set). In addition, our genomic analysis was focused on the molecular profile of GC, particularly involving alterations of EMT-genes, given their role in cancer progression as epithelial cell transformation to mesenchymal cells is fundamental to metastasis (32) and chemoresistance (33,34). Our results coincide with those previously reported in the literature, which provides validity and robustness to our findings and allows us to report novel data or not yet explored potential diagnostic, prognostic, and treatment response markers.

Globally, the alteration profile in GC was dominated by gains. This phenomenon, where gains are more abundant than losses, has been previously reported in different tumor lines, including gastric cell lines (35). Chromosomal gains in cancer might result in increased gene functions, providing cancer cells a competitive advantage for the development of metastasis (36), while chromosomal losses might involve the down-regulation of tumor suppressor genes (37), disrupting homeostasis and accelerating cancer progression. The most affected CNA-chromosomes for DGC were 1, 4, and 5; for IGC 1, 8, and X, and for NAG 6, 7, 14, 17, and X. The altered cytobands associated with GC found in this work (Table 3) are in agreement with previous studies; as an example, 8q24 has been implicated in the development of different tumors (38). The highest frequencies of gains in advanced GC were found at 8q24.21 (65%) and 8q24.3 (60%), and the pattern of CNA in advanced GC was quite different from that in early GC, this increased CNA numbers is associated with disease progression from early to advanced GC (39). The 8q24 cytoband has also been reported in Latin American countries such as Brazil (40) and Venezuela (41), as well as in Asian countries like Korea (42).

Interestingly, the most frequent CNA length in GC was 100-200 Kb in both DGC and IGC *versus* 1-50 Kb in NAG. The biological implications of this alteration length pattern in GC compared to non-cancerous tissues such as NAG is yet to be determined. Meanwhile, it is important to highlight that a resolution of 100-200 Kb *versus* Mb is an advantage of molecular resolution approaches over classical cytogenetics (CGH, FISH, among others) to discover "small" potentially important alterations in cancer samples.

The cumulative length averages (Mb-cl) of these alterations were 183.44 for DGC, 113.56 for IGC, and 1.19 for NAG. These lengths, whether gained or lost, describe the magnitude of global alterations per tissue; yet, their relevance lies on the molecular functions, biological process, and interaction networks in which they participate.

We identified a molecular profile that distinguishes GC from NAG, based on 2441 genes affected by CNA. They are associated with GC, as well as the differences and similarities between histological subtypes, DGC (undifferentiated) and IGC (well-differentiated) compared to a non-cancerous tissue such as NAG (43). Interestingly, we identified 60 affected genes shared between GC and NAG, 19 were shared exclusively with DGC while only eight with IGC. This emerging pattern of shared altered genes between cancerous and non-cancerous tissues, should be further studied to identify possible CNA-dependent oncogenic pathways and progression trajectories from NAG to either GC subtype, particularly in conjunction with environmental factors such as *H. pylori* infection, diet, and lifestyle, that might be involved in spread patterns affecting patient survival (43).

In the heatmap (Figure 3B) a separation between NAG and GC is observed, showing clusters based on the molecular profiles of CNA-genes. There is a greater heterogeneity between the IGC samples in clusters, but there are more genes affected in DGC. The front-line tool for IGC distinction has been based on different criteria such as the histopathological classification proposed by Lauren. However, due to the challenges, conflicts in the correct assignment, diagnosis and treatment new criteria have been proposed such as the molecular characterization by The Cancer Genome Atlas Research Network (TCGA), which divides GC into four subtypes (44). Our observations agree with the need for new proposals for the classification of GC, which includes defined subgroups with the integration of several genomic and genetic parameters where CNA are present.

We analyzed and determined the molecular function profile of GC CNA-genes. Concerning gains, we observed increased alterations involving transcription, signaling, tyrosine kinases, growth factors, hormones, insulin; while in losses, molecules involved in transcription, serine/threonine and MAP kinases, hormones, steroids, beta-catenin binding, filament binding were decreased. These gene sets are important in GC biology. CNA-IGC genes were 13, e.g. CDH1, LAST1, ROCK1, and WWOX; and CNA-DGC genes were 49, e.g. CRIM1, EGFR, MIR9-1, MUC1, MYC, NDRG1, SCRIB, SNAI2, VEGF, and ZEB2, hence, we focus on these example genes with the intention of comparing our findings with others and placing them in a biologically coherent context. For example, CDH1 codes for E-cadherin and, from a simplified viewpoint, E-cadherin maintains the epithelial phenotype; if CDH1 is lost, this promotes the mesenchymal phenotype, i.e., it favors loss of adhesion and metastasis (32).

We performed an enrichment analysis of unique CNA-genes for all tissues and observed several shared molecular functions, such as protein binding. Some gained genes coding for RNA-binding proteins (RBP) (45) have diverse targets and participate in tumor progression by regulating homeostasis and changing expression patterns. Chromatin binding is another altered function in GC that participates in regulating eukaryotic gene expression, methylation profiles modulation, and genome stability maintenance (46). EMT is a process that involves changes in histone modification, DNA methylation, and chromatin accessibility. These changes can be promoted through transcription, allowing the cell to have an identity or to have a MET-EMT conversion (47). The kinase function in DGC and IGC gains (48) have recently been considered key regulators in the development of cancer. Many kinases are related to the initiation and progression of carcinogenesis, and are one of the main therapeutic targets for the development of inhibitors in the clinical area. Kinases are able to promote EMT and enhance invasion, migration, and evasion of apoptosis (49). In IGC, we highlight PIK3R1 and PIK3CA. The PI3K pathway is a key regulatory hub for cell growth, survival and metabolism (50). Activation of PI3K is a frequent hallmark of cancer, highlighted by the prevalence of somatic mutations in genes encoding key components of this pathway (51). These enzymes are responsible for transferring a phosphate group; however, the reverse process is carried out by phosphatases which also are particularly affected in IGC. PIK3R1, is a gene frequently affected by mutations or copy numbers in various types of cancer according to the TCGA project. These genes converge on the PI3K/AKT/mTOR pathway, involved in the regulation of many processes (51). At present, the differences between DGC and IGC have been insufficiently explored and understood; differences in etiology, location, incidence, genetic profiles, among others, have been observed (52). The GC CNA-EMT network (Figure 5) was generated with relevant genes according to different criteria: frequency among patients as well as genetic connections, reported pathways, and experimental associations with several databases: EMTDB, The Human Protein Atlas, COSMIC (53), Cancer Hallmark Genes (CHG) database (54). Shared and exclusive altered genes were observed for each tissue-type. The common CNA-EMT genes between DGC and IGC include GLI2, associated with proliferation (55); EP300, with multiple functions as an inhibitor of anti-tumor immune response via metabolic modulation (56); PTPN11, associated with GC progression; and NDRG1, associated with metastasis and poor prognosis in GC (57). A relevant gene in DGC is EGFR, which given its consistent CNV-GC association, is now the target for the development of anti-GC therapies (58). IGC-exclusive EMT-genes are MICAL2, MYC, and PIK3R1. MICAL2, a destabilizing F-actin in cytoskeletal dynamics, has been found in poor prognosis GC (59). MYC gains have also been reported in several GC studies, as expected for a common oncogenic gene (60) associated with proliferation, differentiation, and apoptosis (61). PIK3R1 participates in the PI3K/AKT signaling pathway with roles in apoptosis and cell survival, as well as chemotherapy resistance in GC (62).

A large amount of data remains to be analyzed, including LOH, mosaicism, and other gene sets that participate in different hallmarks of cancer. Another limitation of our study was the absence of the transcriptomic exploration to validate our GC EMT-signature, particularly for DGC and IGC. Yet, the concordance of CNA with expression alterations of EMT-related genes is plausible, as previously observed for multiple cancer types from The Cancer Genome Atlas (TCGA) data (35). Also, further inclusion of precancerous stages would allow us to depict the “profile” of IGC progression. The results from our genomic approach, coincide with those already reported in the literature, which gives validity and solidity to our results. After all, this strategy allowed us to report novel data, scarcely or not yet explored, to identify differential GC CNA, associate them to relevant molecular functions related to the hallmarks of cancer, and predict the EMT-signature for DGC and IGC. We believe that targeting these networks will potentially serve as diagnostic, and prognostic markers. Additionally, the use of NAG as a non-malignant control allowed us to study the molecular and cellular events of GC, and identify possible biomarkers for the “early” gastric cancer stages.

## Acknowledgments

The authors are grateful to Irma P. Ramos-Vega, Brian-Alexander Cruz-Ramírez, and Alejandra García-Bejarano for technical assistances.

## Availability of data and materials

All data generated or analyzed during this study are included in this published article. The data have been deposited in the Gene Expression Omnibus (GEO; https://www.ncbi.nlm.nih.gov/geo/) using accession number: GSE117093.

## Authors’ contributions

VLS, HAVS, and JDME: Carried out molecular techniques, participated in the analysis of results, preparation, writing, and discussion of the manuscript. JT, MCP, and PPS: Responsible for the clinical aspects of the study, including the recruitment of patients; participated in data discussion and edition of the manuscript. MERT: Responsible for the design of the study. Supervision of the experimental work. Critical review, editing, and writing of the manuscript. JT and MERT responsible for acquiring financial support. All authors read and approved the final manuscript and agreed to be accountable for all aspects of the work in ensuring that questions related to the accuracy or integrity of any part of the work are appropriately investigated and resolved.

